# Broadly-reactive IgG responses to heterologous H5 prime-boost influenza vaccination are shaped by antigenic relatedness to priming strains

**DOI:** 10.1101/2021.04.09.439262

**Authors:** Jiong Wang, Dongmei Li, Sheldon Perry, Shannon P. Hilchey, Alexander Wiltse, John J. Treanor, Mark Y. Sangster, Martin S. Zand

## Abstract

Prime-boost vaccinations of humans with different H5 strains have generated broadly protective antibody levels. However, the effect of an individual’s H5 exposure history on antibody responses to subsequent H5 vaccination is poorly understood. To investigate this, we analyzed the IgG response to H5 A/Indonesia/5/2005 (Ind05) vaccination in three cohorts: (1) a double primed group that received two H5 vaccinations: A/Vietnam/203/2004 (Vie04) 5 years ago and A/Hong Kong/156/1997 (HK97) 11 years ago, (2) a single primed group that received Vie04 5 years ago, and (3) an H5-naïve group that received two doses of the Ind05 vaccine 28 days apart. Hemagglutinin (HA)-reactive IgG levels were estimated by multiplex assay against an HA panel that included 21 H5 strains and 9 other strains representing H1, H3, H7, and H9 subtypes. Relative HA antibody landscapes were generated to quantitatively analyze the magnitude and breadth of antibody binding after vaccination. We found that short-interval prime-boosting with the Ind05 in the naïve group generated a low anti-H5 response. Both primed groups generated robust antibody responses reactive to a broad range of H5 strains after boosting with Ind05; IgG antibody levels persisted longer in subjects who had been double primed years ago. Notably, the IgG responses were strongest against the first priming H5 strain, that reflecting influenza virus immune imprinting. Finally, the broad anti-H5 IgG response was stronger against strains having a small antigenic distance to the initial priming strain.

**IMPORTANCE:** The antigenic shift and draft of hemagglutinin (HA) in influenza viruses is accepted as one of the major reasons for immune evasion. The analysis of B cell immune responses to influenza infection and vaccination is complicated by the impact of exposure history and antibody cross-reaction between antigenically similar influenza strains. To assist in such analyses, the influenza “antibody landscape” method has been used to analyze and visualize the relationship of antibody mediated immunity to the antigenic distance between influenza strains. In this study, we describe a “relative antibody landscape” method, calculating the antigenic distance between the vaccine influenza strain and other H5 strains, and using this relative antigenic distance to plot with the anti-H5 IgG levels post-vaccination. This new method quantitatively estimates and visualizes the correlation between the humoral response to a particular influenza strain, and the antigenic distance to other strains. Our findings demonstrate the effect of H5 exposure history on H5 vaccine responses quantified by the relative antibody landscape method.

## INTRODUCTION

A number of highly pathogenic avian influenza (HPAI) A viruses, such as the H5, H7, and H9 strains, pose a significant threat to cause human pandemics as a result of their fast mutation rate and high pathogenicity (1, 2). To date, there is no evidence of sustained human-to-human transmission of these strains, despite repeated documentation that humans can contract these viruses from infected poultry (3). The first known human H5N1 infection was reported in 1997 during a poultry H5 outbreak in Hong Kong (4). From 2003 to January 2015, a total of 694 laboratory-confirmed human H5 cases were reported across 16 countries and 58% of those people have died as a result (5). Vaccination against future pandemic strains is the most viable path towards mitigating potential outbreaks. However, current H5 non-adjuvanted monovalent influenza vaccine (MIV) formulations are poorly immunogenic (6, 7, 8, 9, 10), and generally require a prime and boost strategy in order to achieve protective levels of immunity (11, 12). Interestingly, boosting with non-adjuvanted MIV, even in subjects who had been primed several years prior lead to robust and broad antibody responses to variant H5 MIV vaccine(11). Such prime and boost strategies also appear to be needed for recent RNA vaccines(13) to other non-influenza viruses; and understanding the immunobiology of this phenomenon remains highly relevant.

It has been generally accepted that the immunological protection against influenza infection is predominately due to antibodies directed against the viral surface hemagglutinin (HA) protein, which is thus the major target of most influenza vaccines(14). A specific language has evolved to describe the potential confounding effects of such exposure on the development of subsequent immunity to influenza. HA imprinting is the initial exposure to an influenza strain, first described in chilhood H1 influenza, which emerging evidence suggests may protect from subsequent H5 infection (2). However, when a person is sequentially exposed to two related virus strains, they tend to elicit an immune response dominated by antibodies against the first strain they were exposed to(15, 16). This is true even following a secondary infection or vaccination. This phenomenon has been variously referred to as “original antigenic sin” (OAS), HA seniority, or negative antigenic interaction (17, 18, 19).Thus, the immune response to a new influenza viral infection or vaccination is at least partially shaped by preexisting influenza immunity. Because there is still antigenic overlap between even mostly dissimilar influenza strains, it is critical to understand the antibody response against antigenically similar virus stains, for vaccine development, especially within the context to OAS.

The HA protein is composed of two domains, the highly plastic globular HA1 head domain and the conserved HA2 stalk domain. The hypervariable head domain is believed to be immunodominant and virus infection or/and vaccination elicits strain-specific neutralizing antibodies primarily targeting this domain, resulting in limited cross-reactivity to divergent virus strains that vary significantly in HA1 head domain sequence(20). In contrast, antibodies targeting the conserved stalk HA2-reactive domain have been shown to broadly cross-react with multiple influenza viral strains (21). The viruses themselves can be categorized based on the phylogenetic distance of HA sequences. Ten clades of H5 HA (clade 0-9) have been identified within the H5N1 virus subtype (22). H5N1 viruses from clades 0, 1, 2, and 7 have the capacity to infect humans (23). These scatter into three distinct antigenic clusters, as determined by antigenic cartography generated by analyzing neutralizing serum antibody levels elicited in mice vaccinated against single influenza strains (1). As such, an effective H5 influenza vaccine would ideally induce broad cross-reactivity that against all three H5 clades. However, as discussed above, HA imprinting or OAS may impede generation of broadly cross-reactive H5N1 antibodies if the prime and boost H5N1 vaccine strains reside in different antigenic clusters.

To address this issue, we re-analysed serum samples from a previous H5 human vaccine study (DMID 08-0059)(24) using our mPlex-Flu assay multiplex assay(25) to measure the anti-HA IgG antibody against all 10 clades (subclades) of H5 influenza virus. During this study, longitudinal samples were collected prior and post-vaccination with inactivated A/Indonesia/5/05 (Ind05) MIV from subjects: a) who had received two primed H5 MIV vaccinations (A/Hong Kong/156/97 (HK97) in 1997–1998 and A/Vietnam/1203/04 (Vie04) in 2005–2006 (DL-boost group); b) who had only received one Vie04 prime vaccination in 2005-2006 (L-boost group); and c) H5 influenza virus naive group, who were also boosted by Ind05 28 days after the prime event (S-boost group). The mPlex-Flu assay(25) enables us to simultaneously evaluate the magnitude and breadth of the IgG repertoire directed against HAs from 21 H5 influenza virus strains and 9 other IAV strains (H1, H3, H7, H9). We also introduced a novel multiple dimensional data analysis method: relative antibody landscapes, which enables quantitative analysis of the antibody response to influenza virus antigenic similarity strains related to vaccine strains. The relative antibody landscape enables analysis of antibody-mediated immunity to a spectrum of HAs after H5 vaccine priming and boosting. This report demonstrates that as the relative antigenic distance between the original priming and the new H5 boosting vaccine strain becomes smaller (i.e. the strains are more antigenically similar), the greater the increase in the anti-HA IgG response to original H5 MIV strain. Thus, in a vaccine response, the original HA imprinting influences vaccine responses occurring significantly later. We discuss the relevance of these findings to the development of influenza vaccines designed to induce broad antibody-mediated protection.

## RESULTS

### Characteristics of subjects

Prior exposure to the predominant seasonal H1 or H3 influenza strain circulating close to a subject’s birth year can alter H5 or H7 infection and death rates (2, 26). Thus, we first tested tested for differences in age, as a surrogate for circulating strains, that could alter the antibody levels between the H5 vaccine groups. To assess the birth year related influenza virus exposure history, we regrouped the study cohorts based on two key birth years: 1968 and 1977, when H3 and H1, respectively, became the dominant circulating influenza A virus strains (Table 1) (2). Subjects without baseline (pre-vaccination) serum samples were excluded, leaving a total of 55 subjects. The H5 naive subjects (Naive, *n* = 12) and primed subjects (L-boost, *n* = 30) previously received an inactivated subvirion influenza A/Vietnam/1203/04 (Vie04) vaccine in 2005–2006(11). The double primed group (DL-boost, *n* = 13) received the recombinant influenza A/Hong Kong/156/97 vaccine (A/HK97) in 1997 - 1998 (6) and the Vie04 vaccine in 2005 - 2006. We found no significant difference in birth year distributions between the cohorts (P > 0.05; Fisher’s exact test), suggesting that the effects of flu exposure history on the H5 MIV vaccine response should be similar across the three groups.

**TABLE 1.**
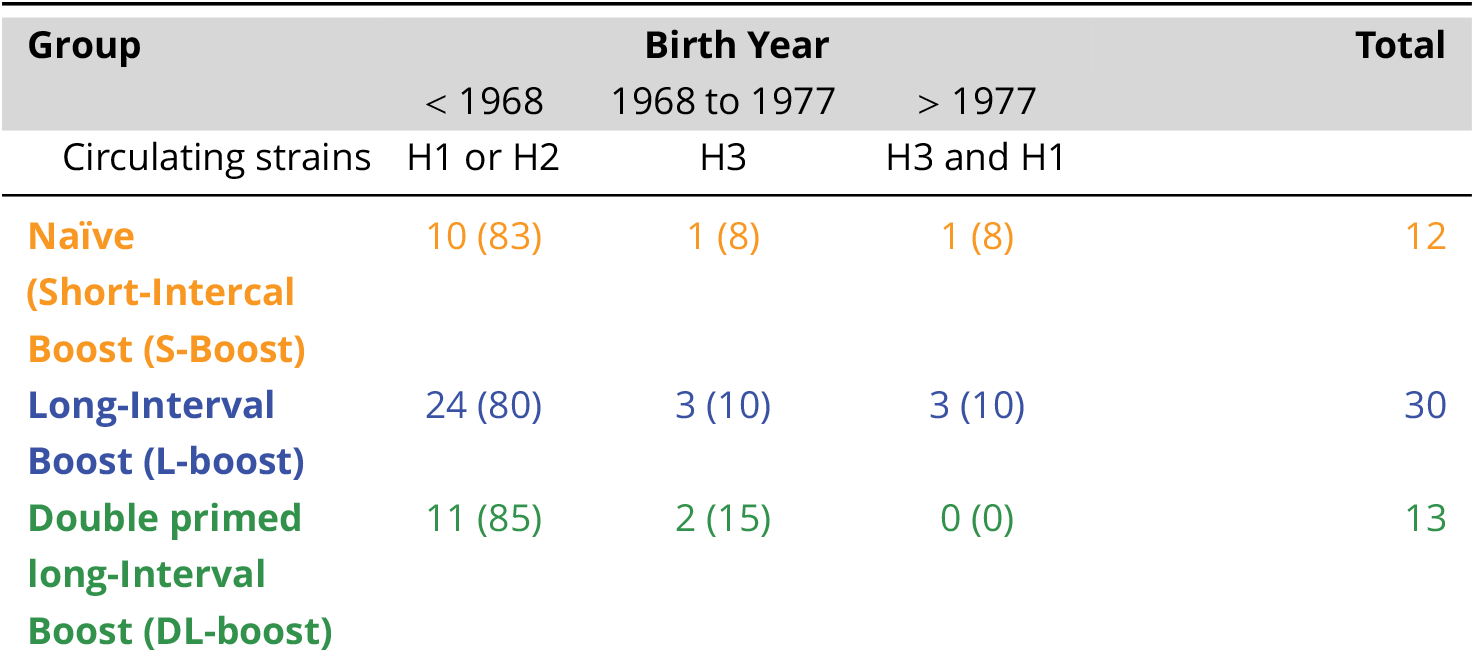
The number of subjects stratified by birth year in each cohort of the DMID 08-0059 study. Subjects were grouped by birth year based on key years when either H3 or H1 representing the predominant circulating seasonal flu strains, as prior exposure history might influence the antibody responses to the H5 vaccines.

### High anti-H5 IgG responses after long-interval boosting are shaped by the priming vaccine strain

Using a 48-HA mPLEX-Flu assay panel, we observed that IgG levels against the HA of A/Indonesia/5/05 (Ind05), Vie04 and HK97 were very low in the naive group, and about two-fold higher in the short interval boosting (S-boost) group who were boosted after 28 days (FIG 2 A, B). In both primed groups (L-boost and DL-boost), however, inactivated Ind05 MIV induced ∼5-fold higher vaccine-specific antibody levels by 14 days post-vaccination. Anti-Vie04 and HK97 IgG levels increased ∼7-8 fold, also peaking at 14 days in both primed groups (FIG 2). While both primed groups had higher pre-existing (day 0) anti-H5 IgG levels, their IgG response kinetic curves against the vaccine strains were similar. These differences result in a relative increase in the DL-boost group’s anti-HA antibody levels peaking at 3.5-fold (FIG2, D), even though the post-boosting IgG levels are similar in the S- and DL-boost groups. In both groups, anti-H5 HA antibodies levels remained high for over six months. These results are consistent with the previous finding that non-adjuvanted MIVs are poorly immunogenic in naive subjects (6, 7, 8, 9, 10), and long-interval boosting with H5 antigenic variant MIVs elicits significant and robust antibody responses (11, 24). However, this is the first report to show differences in antibody response induced by single vs. double long-interval MIV boosting.

**FIG 1.**
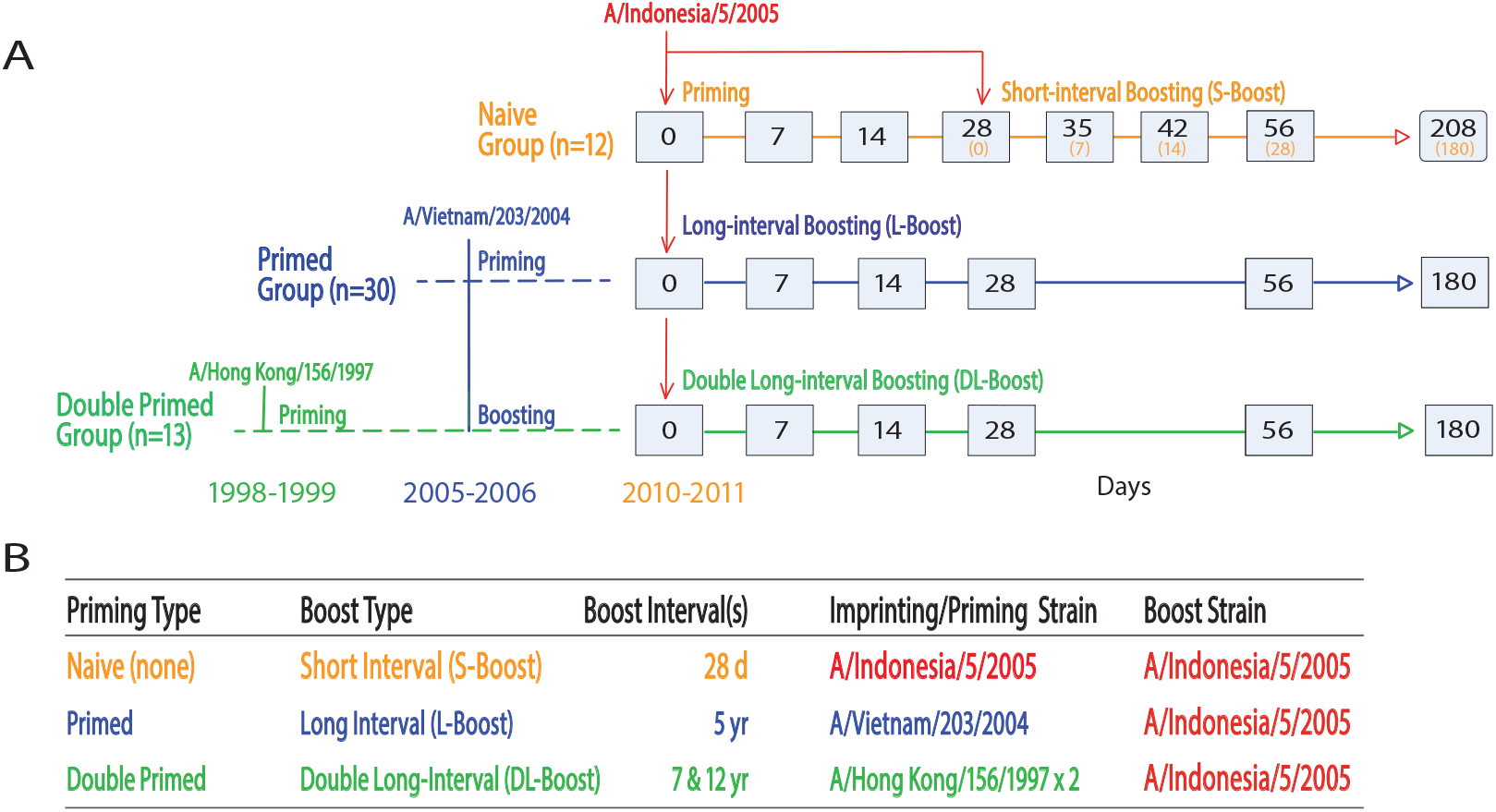
**Vaccination strategy. (A) Trial and sampling design: All subjects in the DMID 08-0059 study cohorts were vaccinated with inactivated A/Indonesia/5/05 (Ind05) intramuscular influenza vaccine. The Naive group; S-boost received the Ind05 vaccine on day 0, and short interval boosting on day 28. The primed long-interval boost (L-boost) group had previously received the inactivated subvirion influenza A/Vietnam/1203/04 (Vie04) vaccine in 2005–2006; and the double primed long interval boost (DL-boost) group additionally received the baculovirus expressed recombinant influenza A/Hong Kong/156/97 vaccine (HK97) in 1997–1998. Both L-boost and DL-boost groups also received long-interval vaccination with Ind05 on the day 0. Grey boxes indicate serum sampling. A) Summary of prime and boost strains and groups.**

**FIG 2.**
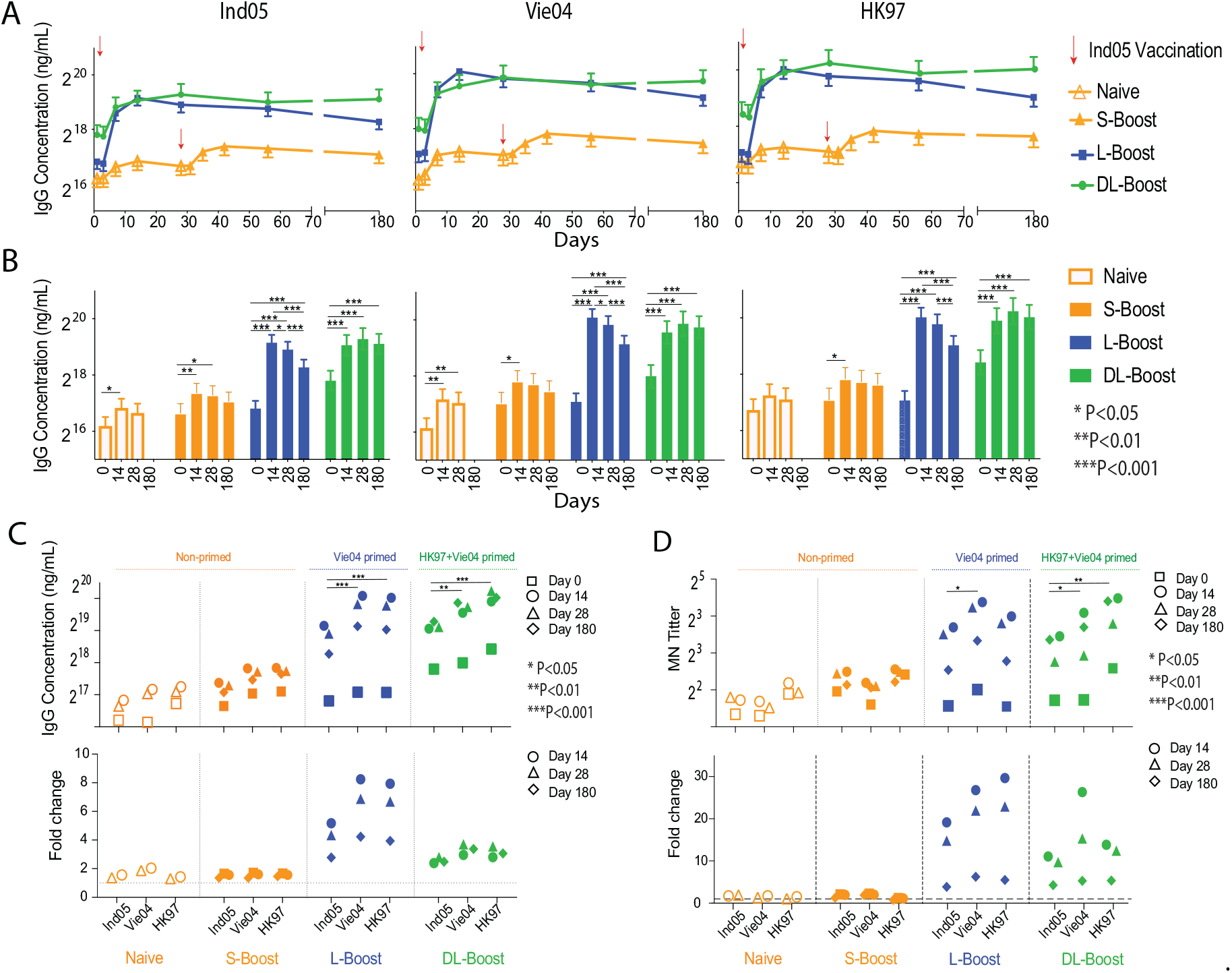
**The effects of prior vaccination with H5 monovalent influenza vaccine (MIV) on multiplex HA antibody responses against three different H5 virus boosting vaccine strains. The mean and standard deviation of IgG concentration for each group were estimated by the mPlex-Flu assay. Antibody concentrations were adjusted within the linear mixed effects models using age at enrollment, gender, ethnicity (Caucasian vs. non-Caucasian), dose (two dose levels: 15 and 90 *µg*), and assay batch (five batches) (27, 28). A. The H5 kinetic antibody levels against three vaccine strains after MIV H5 vaccination with A/Indonesia/05/2005 (Ind05; clade 2). The prime response (Naive, unfilled symbols) and short-interval boost response (S-boost, filled symbols) of naive subjects; the long-interval boost response (L-boost) after one dose of Ind05 MIV in subjects primed by Vie04 MIV the 5 years previously, and in the subjects who were double primed with Vie04 (5 years previously) and A/Hong Kong/156/1997 (HK97; clade 0) HK97 (12-13 years previously), as the double long-interval boost response (DL-boost), against Ind05, A/Vietnam/1203/2004 (Vie04; clade 1) and A/Hong Kong/156/1997 (HK97; clade 0), three vaccine H5 strains. B. Comparison of antibody responses between time points in the same groups for each vaccine strain. C. The antibody concentrations against each vaccination strain and fold changes as compared to day 0, grouped by study cohort. D. The antibody titers for micro-neutralization (MN) against each vaccination strain and fold changes, as compared to day 0, grouped by study cohort. The original MN assay data was been re-analysed with linear mixed effects modeling, as above. Shown are the geometric mean of titers. * P<0.05, **P<0.01, ***P<0.001 Linear contrasts within the linear mixed effects model framework were used to conduct the statistical comparisons.**

Importantly, we also found that the Ind05 MIV elicited robust antibody responses against the two previous priming H5 strains (Vie04, HK97) in both vaccine groups, and that the anti-HA IgG responses shared similar kinetic patterns. Interestingly, Ind05 MIV elicited higher levels of IgG antibodies to Vie04 and HK97 than to Ind05. In order to directly compare the effects of the priming virus strain, we plotted the concentrations of anti-H5 HA by groups, shown in FIG2 C, and the fold change of antibody concentrations against three vaccine strains of the different groups (FIG2 D). The results revealed higher antibody levels against the HA of Vie04 in the L-boost group, and HK97 in the DL-boost group, which were the first H5 viral strains subjects were respectively vaccinated against. These results could be interpreted as indicative of HA imprinting (16, 15), in which subjects generate a robust antibody response against the H5 influenza virus strain they were first exposed to, by infection or vaccination, and maintain this response over their entire lifetime (29).

To confirm the protective activities of the higher level of long-lasting antibodies in the L-boost and DL-boost groups we re-analyzed the HAI and MN data from the DMID 08-0059 study using generalized linear mixed effects models with identity link functions, as we have previously described (27, 28). The results confirmed that all three H5 MIV strain vaccines induced serum with viral neutralizing capacity that could protect cells from viral infection (FIG 2 D and FIG S8).

### Relative antigenic response landscapes of H5 MIV Has

Our results also raised another fundamental question: Does the magnitude of the imprinted recall response to the primoriginal H5 HA correlate with the antigenic distance between the HAs of the prime and boost strains? We hypothesized that the antigenic distance between the vaccine strain and a target H5 HA is inversely correlated with the cross-relativity of antibody response induced by the H5 MIV. In other words, smaller antigenic distances from the first influenza virus strain (imprinting strain) produce larger IgG responses. To answer this question, we performed antigenic cartography to quantitatively evaluate the antigenic distances between H5 clades and subclades.

Recombinant H5 HA proteins were expressed and purified. Strains were chosen to cover all 10 H5 clades (0-9) and subclades, and 4 new H5 avian strains (Cl4.4.4.3) isolated in the US (TABLE S1, and FIG S1). Antibody reactivity to these strains was plotted against mouse anti-H5 HA IgG serum reactivity generated utilizing a monovalent DNA vaccination approach (FIG S2 A). We thus generated a comprehensive antigenic distance matrix between 17 H5 influenza virus strains and each of 21 H5 and 9 other influenza virus strains using the mPlex-Flu assay. The individual antibody levels against H5 viruses are shown as MFI units at specific dilutions, with the dilution factors being normalized using a generalized linear model with an identity link function for the sera samples. We used classical multidimensional scaling (MDS)(30) to project relative distances between strains into 2 dimensions, and the matrix data was created by calculating a Euclidean distance matrix from two-dimensional coordinates. Finally, we used a modification of the approach of Smith, et al.(31) to visualize the antigenic distance between influenza virus HAs(31, 1) (FIG S2, C). This approach accounts for the continuous nature of the mPlex-Flu assay data and the consistent range of estimated strain-specific binding(27, 28), yielding the same results as antigenic cartography. The antigenic distance matrix was also generated from the above multiplex data of mPlex-Flu assay using the single virus DNA vaccine anti-sera. (FIG S3).

In order to show the relative antigenic distance between individual HAs and the H5 MIV strains (FIG 3 B), we plotted the distance of each H5 HA relative to the 3 vaccine strains: HK97 (X-axis), Vie04 (Y-axis) and Ind05 (Z-axis). Each marker diameter represents the magnitude of the IgG concentration 14 days after MIV boosting. This allowed visualization of the magnitude of the antibody response against specific H5 HAs, associated with the antigenic distances with respect to both prime and boost vaccine strains in the different cohort groups. The same diagram allowed visualization of H5 strain vaccine strain relative distances from other H5 strains. Naive subjects had low anti-HA IgG levels against all H5 strains after priming and short-interval boosting with MIV. However, the L-boost and DL-boost groups had significantly enhanced antibody responses after 14 days, with higher IgG responses to H5 strains in the Vie04 and HK97 cluster groups than to the viruses in the MIV Ind05 cluster group, which are antigenically similar to the strain of the more recent MIV (FIG 3 C). These data more clearly show the relationship between the anti-HA IgG antibody response and the antigenic distances to the reference strains: higher cross-reactive antibody levels are elicited against the HAs from strains in the same cluster group with the first priming virus strain.

**FIG 3.**
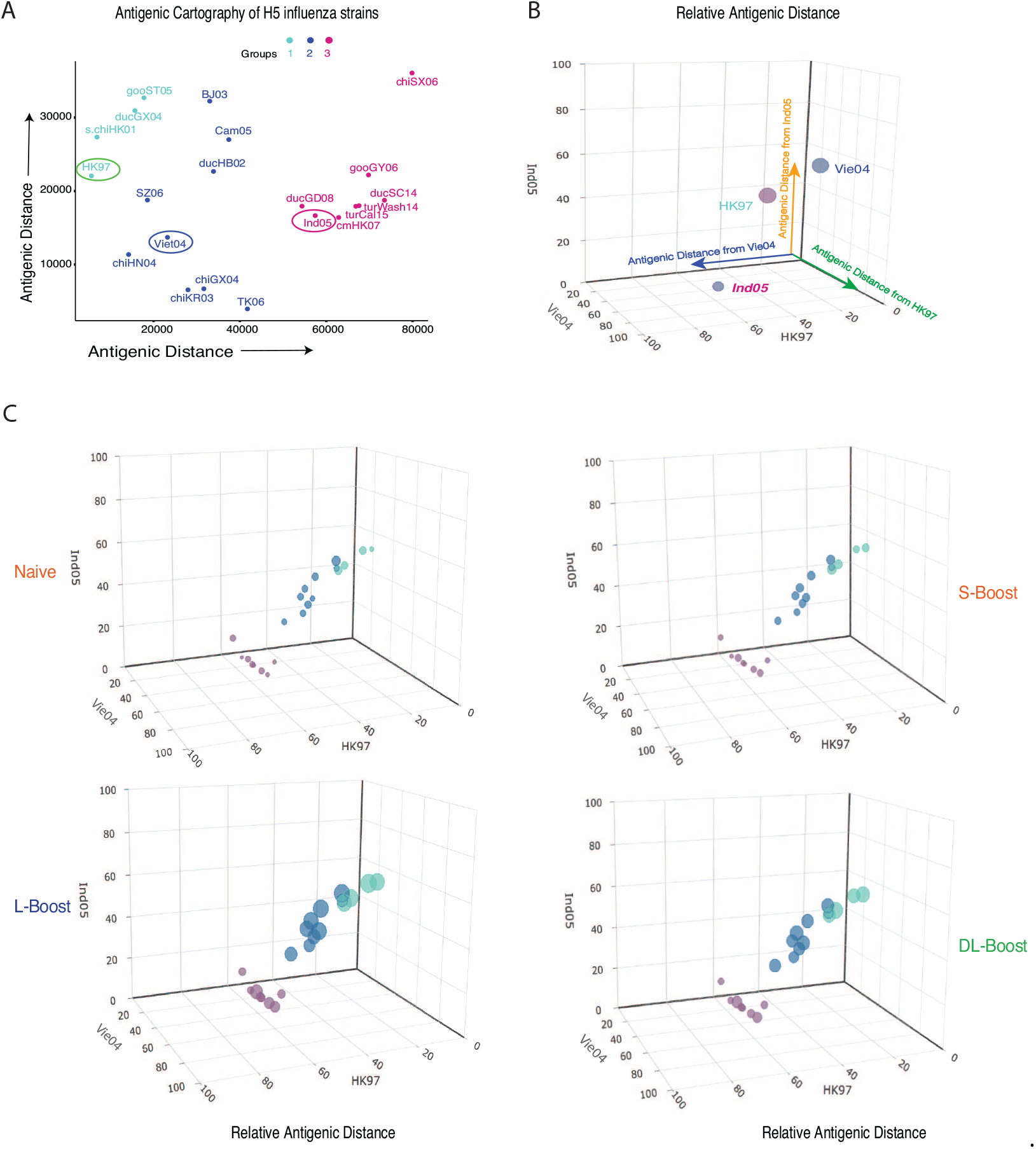
**HA antibody responses were plotted against the related antigenic distances to each monovalent H5 vaccination (MIV) strain of different study cohorts. A. Antigenic cartography of 21 H5 influenza virus strains generated by mPlex-Flu assay of antisera against 17 anti-H5 influenza viruses, and plotted using using classical multi-dimensional scaling (MDS;see Methods). The three vaccine strains are circled. B. We then used three dimensional plots to show the relative antigenic distance of all mPLEX-Flu target HAs to the three vaccine strains A/Hong Kong/97 (HK97, clade 0), A/Vietnam/1203/2004 (Vie04; clade 1), A/Indonesia/05/2005 (Ind05; clade 2). C. The IgG response of subjects in the DMID 08-0059 study to 21 H5 strains plotted in 3D bubble plots. The relative antigenic distances of the 21 H5 strains assayed were plotted against their antigentic distance to each of the three MIV strains to determine giving 3D-antigenic cartography. The bubble size represents the concentration (10^4^ng/mL) of IgG against an H5 influenza virus at day 14 post MIV boosting. (A) Using unsupervised hierarchical clustering, three H5 antigenic groups were identified. Interactive 3D bubble plots can be accessed through the following links: Prime group (http://rpubs.com/DongmeiLi/565996); S-boost group(http://rpubs.com/DongmeiLi/565998); L-boost: (http://rpubs.com/DongmeiLi/565989); DL-boost: (http://rpubs.com/DongmeiLi/565994).**

### Long-interval boosting (L-boost) of MIV elicited heterogeneous IgG responses against all H5 clade/subclades, which were correlated with the antigenic distance to the first primed virus strains

We next generated antigenic landscape plots (26) to visualize the magnitude of serological responses in relation to the antigenic distance between the vaccine strain HA and the H5 HAs in the mPlex-Flu panel. We first focused on the relationship between the magnitude of boosted IgG response and the antigenic distance between the boost HA and the three H5 vaccine strains. To this end, IgG antibody concentrations against 21 H5 strains were measured by mPLEx-Flu assay for each cohort on days 9, 14, and 28, which were plotted against their relative antigenic distances to Ind05 (FIG 4A, B), Viet04 (FIG 4C, D), and HK97 (FIG 4E, F). Correlation test results are given in the figure inset, and all data are presented in FIG S4, S5, S6).

**FIG 4.**
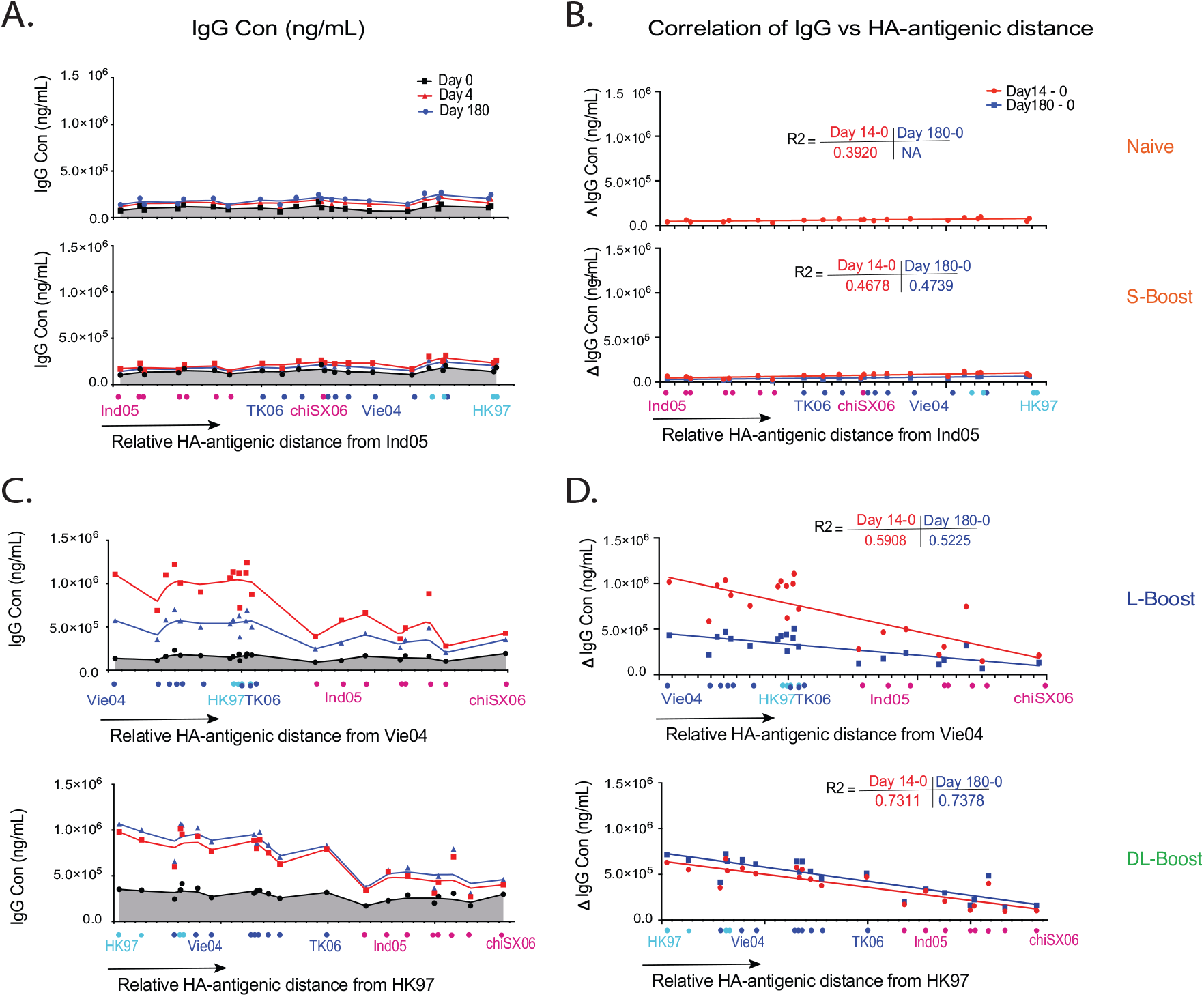
**Relative HA antibody landscapes, anti-HA IgG levels and relative antigenic distances from vaccine strains. A. The relative HA antibody landscapes of H5 virus strains as a function of the relative HA antigenic similarity distance from the vaccination strain Ind05 for the Naive group and short interval boost (S-boost) group (see Materials and Methods). B. Correlation of the HA antibody response to the HA-antigenic distance from the vaccine strain HAs of the Naive and S-boost groups. The coordinates of each H5 strain result represent the relative antigenic distance of H5 HA_*i*_ to the vaccine strain HA on each axis. C. Relative HA antibody landscapes for each group using the relative HA antigenic distance from the H5 reference strains A/Vietnam/1203/2004 (Vie04; clade 1), or A/Hong Kong/97 (HK97, clade 0). D. The correlation between the HA antibody response and the HA-antigenic distance to the imprinting (first exposure) H5 strain: Vie04 for the long-interval boost group (L-boost)or HK97 for the double long-interval boost group (DL-boost). The change of IgG concentration (Δ*IgG*_*conc*_) is the difference between the anti-HA antibody concentration of past-vaccination from that of prior vaccination. The *R*^2^ values were calculated from linear regression fitting.**

We found that the immune response in the Naive and S-boost groups were very weak, and since subjects in these groups were only exposed to the Ind05 MIV strain, we made antigenic landscapes (26) using Ind05 as the reference influenza virus strain. The relative antigenic landscapes for these two groups at days 0, 14 and 180 are shown in FIG4 A and B. Similarly, the serological responses of the L-boost and D-boost groups after boosting were plotted against the antigenic distance relative to Vie04 and HK97, shown in FIG4 C and D. Note that the antigenic distance between the cognate vaccine strain and itself is zero (e.g. Vie04 - Vie04 = 0). The Ind05 MIV showed very low antigenicity in both naive subject groups. Changes in IgG concentration (Δ*IgG* = [*IgG*_*t*_] − [*IgG*_*day*0_]) were not correlated with antigenic distance (P = 0.014 and 0.020). However, Ind05 MIV boosting showed higher antibody responses to HAs from strains with a smaller antigenic distance in both L-boost (*R*^2^ = 0.57) and DL-boost groups (*R*^2^ = 0.73). These results support our hypothesis that that the imprinting of primed individuals is highly correlated with the related antigenic distance to the priming strains for long-interval H5 vaccination. FIG 4.

### Long-interval boosting with H5 MIV induced broadly heterosubtypic antibody responses against Group 1 influenza viruses

To assess the breadth of heterosubtypic immunity generated by the H5 MIV prime and boost strategy, including IgG reactive against other influenza virus HAs, we estimated antibody cross-reactivity to select group 1 (H1, H2, H5, H6, and H9) and group 2 (H3, H4, H7) HAs (Table S1)) using the mPlex-Flu assay (FIG 5). In all subjects, we detected high pre-existing anti-H1 HA subtype IgG levels against older (A/South Carolina/1/18 (SC18), A/Puerto Rico/8/1934 (PR8)) and newer (A/New Caledonia/20/1999 (NewCall99), A/California/07/2009 (Cali09)) strains. However, these anti-HA levels were not significantly affected by H5 MIV vaccination (FIG S7 A). In addition, we found dramatic increases in anti-HA IgG levels targeting other group 1 influenza viruses (e.g. H2, H6) that had lower baseline levels compared to those against influenza group 2 (H1, H3) subtype virus HAs.

**FIG 5.**
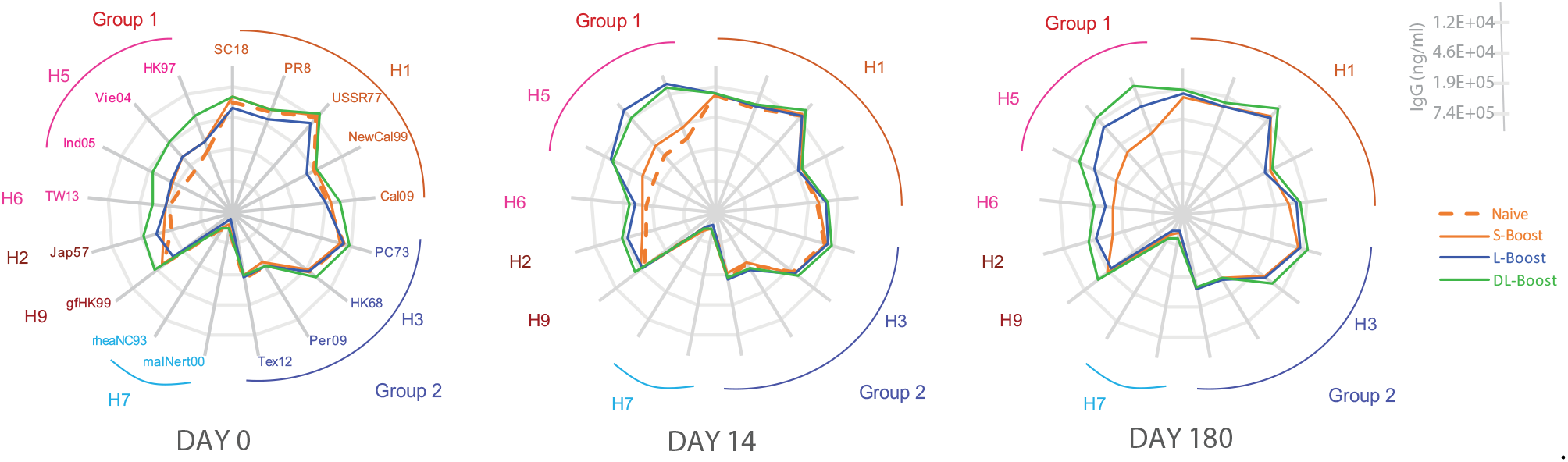
**The human heterosubtypic IgG antibody response elicited by H5 MIV. The IgG antibody response induced by H5 influenza vaccine against previously circulating or vaccine virus strains (H1, H2, H3, H6, H7, H9), were measured by mPlex-Flu assay pre-(day 0) and post-vaccination (days 14, 180).**

Further analysis demonstrated that post-H5 vaccination IgG reactivity across influenza virus strains was inversely correlated to both phylogenetic and antigenic distance between the strains, especially the stalk regions. Based on phylogenetic distance, the gene sequence of H6 is closer to H5 than H9 (20). Similarly, the gene sequence of H2 is closer to H5 than H6 and H1 (FIG S1 A). In addition, we found that IgG responses induced by H5 MIV against HA of A/Japan/305/1957 (Jap57, H2) were significantly higher than that against A/Taiwan/2/2013 (TW13, H6) and A/guinea fowl/Hong Kong/WF10/1999 (gfHK99, H9) (FIG5, FIG S7 A), the latter two strains have stalk regions phylogenetically and antigenically distant from the H5 clade stalk. We also found that, in both primed groups, H5 MIV elicited cross-reactive anti-H2 IgG responses in naive subjects, with a higher peak and a sustained duration than in the Naive subjects. Those responses were stronger than those against H6 and H9 HAs. No significant changes were detected in IgG levels against H3 and other group 2 influenza viruses (FIG S7 B). Together, these findings also support the hypothesis that cross-strain, anti-HA antibody responses are highly correlated with phylogenetic similarity, and inversely correlated with antigenic distance, to the vaccine strain.

### Long-interval boosting elicited IgG antibodies against the HA head domain

The HA stalk domain is highly conserved within influenza virus phylogenetic groups, and stalk-reactive antibodies have been hypothesized to be the major contributors mediating cross-reactivity of anti-HA IgG antibodies across group 1(32) strains. However, broadly cross-reactive neutralizing antibodies against the HA head domain have recently been identified, and could also contribute to this phenomenon (reviewed in (33)). Thus, we next measured the change in the relative proportions of head versus stalk reactive IgG within H5 boosting group.

H5 head (HA1) specific IgG levels were measured using beads coupled with the Ind05 head domain only. Anti-stalk IgG was measured using chimeric cH9/1 and cH4/7 proteins to estimate, respectively, group 1 and group 2 stalk-reactive antibodies (34, 35, 36). The results demonstrate that short-interval boosting can induce an ∼2 fold increase in anti-H5 head IgG levels in naive subjects (FIG 6). In addition, significant increases in head-specific IgG were also detected in the L-boost group: 27 fold (14d), 20 fold (28d), and 10 fold (180d). Examining the DL-boost group, ∼7-8 fold increases were observed at 14, 28, 180 days after vaccination. High levels of group 1 stalk-reactive IgG were found in both boosting groups. However, these increases accounted for less than a 2-fold overall change in IgG levels, primarily because these stalk-reactive IgG antibodies were present at relatively high levels prior to vaccination. We did not observe any significant post-vaccination increases in group 2 stalk-reactive antibody levels regardless of test groups. Overall, our results suggest that broadly cross-reactive IgG against H5 influenza virus HAs or the phylogenetic group 1 are most likely mediated by conserved epitopes on the head domain of HA as opposed to the stalk domain.

**FIG 6.**
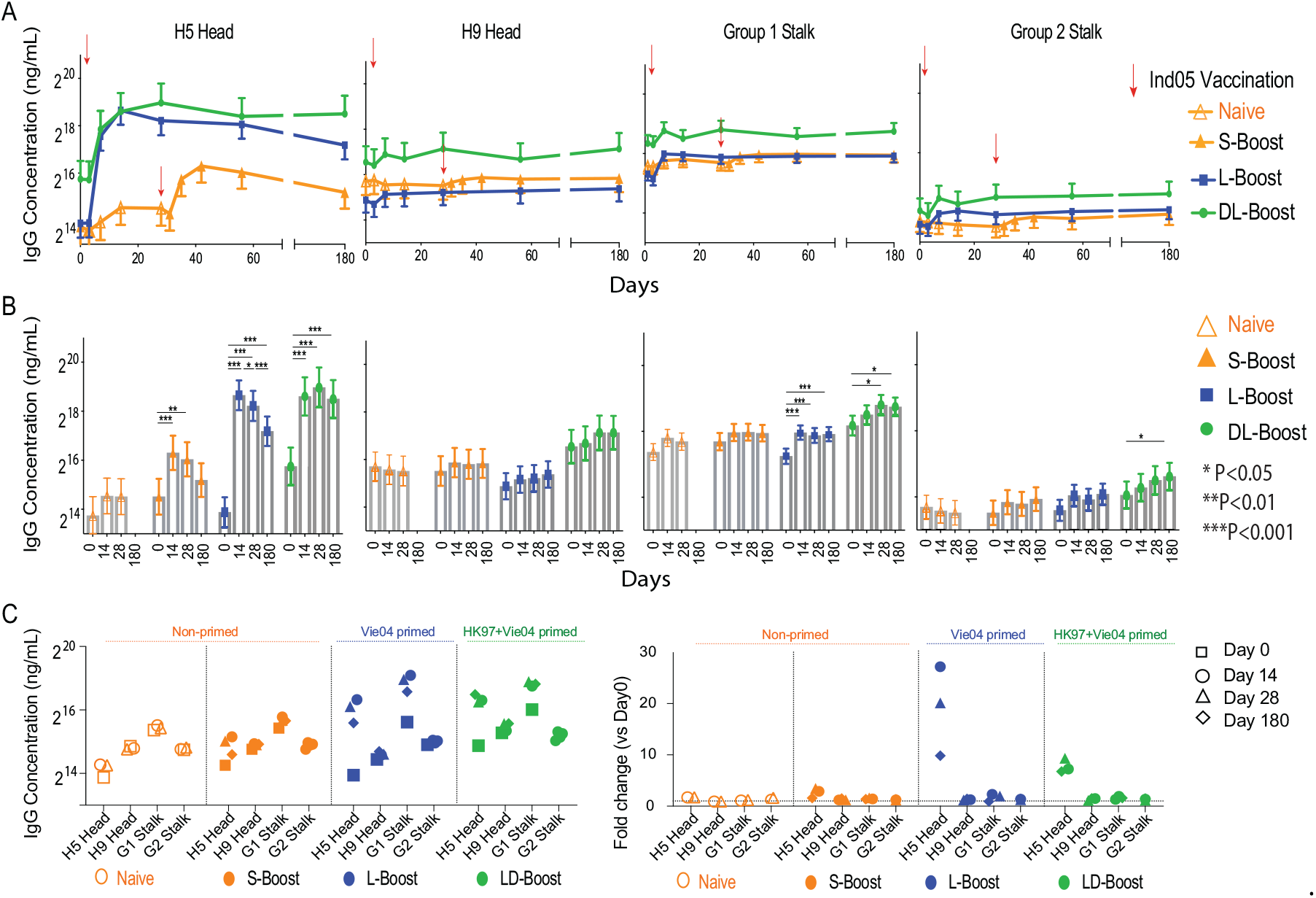
**The head and stalk-reactive IgG response induced by the human MIV H5 vaccine. A. The kinetic profile of the IgG response against the HA head or stalk domain estimated by mPlex-Flu assay. B. Comparison of concentrations of each H5 HA specific antibody pre-(day 0) and post-vaccination (14, 28 and 180 days). Linear contrasts within the linear mixed effects models framework were used for statistic testing (* P<0.05, **P<0.01, ***P<0.001). C. Comparison of anti-HA IgG concentrations between HAs, including antibodies against chimeric cH9/1 HA (termed group 1 stalk-reactive antibodies; G1 Stalk), and cH4/7 HA (termed group 2 stalk-reactive antibodies; G2 Stalk).**

## DISCUSSION

Two major impediments to universal flu vaccine development are the constant antigenic changes of influenza viruses, and that the human antibody response is shaped by prior influenza virus exposure history (37). In addition, vaccination strategies for emergent influenza viruses need to take into account both the vaccination schedule, and the ability of HA imprinting to can hinder immune responses to new antigens. Antibody mediated immune responses to new HA antigens are generally weak after the priming vaccination, and require further boosting to elicit adequate titers for infection prevention. This phenomenon can be leveraged if the subject has been primed by exposure to HA antigens, by prior infection or vaccination of H1 or H3 influenza virus, that are antigenically distance from emergent strain HAs (heterosubtypic immunity).

The antigenic distance between two virus strain HAs can be calculated empirically or experimentally. Empirically, antigenic distance is correlated with the difference between surface protein sequences of HA (e.g. edit distance, Damerau-Levenshtein distance). Experimentally it can be derived by calculating the n-dimensional distance between immune reactivity of sera from a subject vaccinated with a single virus against a panel of other HAs from disparate virus strains (37). As we have previously shown (35), the smaller the antigenic distance between the prime and boost HAs, the stronger the post-boost vaccination increase in vaccine specific anti-HA IgG levels.

In this study, we also analyzed changes in multi-dimensional anti-H5 HA IgG responses after vaccination and boosting using a modification of the antibody landscape method (29), a variant of antigenic cartography (31). We initially analyzed anti-HA IgG antibody levels against a comprehensive panel of H5 clade/subclade HAs as a function of the relative antigenic distance to the reference vaccine HA. We call this multi-dimensional measure the *relative antibody landscape* (Fig 4 A and C). This novel method, combined with multiplex serum IgG measurements, allows an analysis of the breadth of the antibody response as a function of the antigenic distance from the vaccine strain. Our results using the relative antibody landscape method show that the anti-H5 HA IgG responses elicited by boosting in both primed groups are highly correlated with the antigenic distance between the priming and boosting H5 vaccine strains. These findings provide further evidence of for the HA antigenic imprinting in H5 influenza vaccination. Most significantly, we demonstrate that relative antibody landscape methods can be used to analyze the effects of previous HA antigen exposure on vaccine responses, allowing for quantitative analysis of antigenic imprinting.

Our work also demonstrates that long-interval boosting augments H5 vaccine-induced immunity. Studies using variants of the H5MIVs have shown that long-interval prime-boost strategies, on the order of 4-8 years between vaccinations, result in robust and durable antibody responses (11) to what are relatively poorly immunogenic vaccine components (6, 7, 24). Intermediate intervals of 6-12 months between priming and boosting with H5 variants significantly increases antibody responses (38, 39), compared to 8 weeks or less. One potential mechanism for these results is a time-dependent increase in long-lived memory B cells, which may take 2-4 months after vaccine priming (40). These memory B cells can then respond rapidly to long interval boosting (41). Studies showed that adjuvanted H5 MIV used in short-interval boosting also significantly increased the immunogenicity of vaccines (42, 43, 44, 45), and indicated that prime-boost vaccination induced the monoclonal antibodies largely recognized the HA head region of the H5 MIV strain(46). Significant additional work is necessary to define the optimum prime-boost interval for robust responses.

Our results also support the hypothesis that long-interval boosting increases antibody responses targeting the HA head domain, rather than the stalk. Recently, several broadly neutralizing antibodies (bnAbs) have been identified from both infected or vaccinated human subjects that target the hypervariable HA head domain, including C05 (47), 5J8 (48), CH65 (49) and CH67. These bnAbs exhibit considerable neutralizing breadth within the H1 (47, 48, 49) and H3 (50) influenza virus subtypes. Such bnAbs are thought to bind highly conserved regions on the sialic acid receptor binding site (RBS) in the HA head domain, explaining their ability to broadly neutralize viral binding from different subtypes (49, 51). As the head domain is known to be immunodominant in the induction of strong antibody responses, broadly head-reactive antibodies could be the major mediator of cross-reactive immunity across subtypes or heterosubtypes. Our results are also consistent with recent work that found rapid activation and expansion of pre-existing memory B cell responses to the conserved epitopes on the HA stalk and head domains after long interval prime-boost vaccination with H7N9 (40).

Finally, our results contribute further to a framework for thinking about influenza vaccine development strategies. The aspirational goal of a influenza vaccine is to create long-lasting protective immunity to a wide spectrum of influenza viruses. In such cases, future exposure, via infection or vaccination may occur years after the initial priming and imprinting event. Our work demonstrates that the long interval prime-boost strategy for H5 vaccination induces long-lasting cross-reactive antibodies against conserved regions on the HA1 head domain. This may help in universal influenza vaccine development not as a single vaccine, but as a long-interval boost strategy to generate cross-reactive antibodies to recognize the conserved sites on HA1 head domain.

In conclusion, we used a multiplex antibody assay and a novel antibody landscape method to analyze antibody mediated immunity to various HAs after H5 vaccine priming and boosting. These methods quantitatively account for the antigenic distances between the vaccine and other strain HAs. This new approach demonstrated that anti-H5 IgG antibody responses elicited by boosting are highly correlated to the antigenic similarity between the priming and boosting H5 vaccine strains, providing evidence for OAS and HA imprinting within the context of H5 vaccination.

## MATERIALS AND METHODS

### Human Subjects Ethics Statement

This sub-analysis study was approved by the Research Subjects Review Board at the University of Rochester Medical Center (RSRB approval number RSRB00012232). Samples were analyzed under secondary use consent obtained previously as part of prior clinical trial (24). All research data were coded by sample IDs in compliance with the Department of Health and Human Services’ Regulations for the Protection of Human Subjects (45 CFR 46.101(b)(4)).

### Samples and data

Serum samples for the multiplex assay were obtained from a prior clinical trial, DMID 08-0059 (Figure 1)(24). Subjects without pre-vaccination serum samples (Day 0 baseline) were excluded. All subjects in the three cohorts were inoculated with inactivated A/Indonesia/5/05 (A/Ind05) vaccine. H5 naive subjects (*n* = 12), who were healthy adults, not at risk for H5 exposure and with no H5 vaccination history, received 2 identical A/Ind05 vaccinations separated by 28 days. Primed subjects (*n* = 30) previously received the inactivated subvirion A/Vietnam/1203/04 (A/Vie04) vaccine in 2005–2006 (11). The double primed group (*n* = 13) had received both the recombinant A/Hong Kong/156/97 vaccine (A/HK97) in 1997-1998 (6) and the influenza A/Vie04 vaccine in 2005-2006. Serum samples were collected before vaccination (Day 0) and on days 7, 14, 28, 56, and 180 after vaccination. Serum samples were collected from the naive group subjects on days 7, 14, and 28 days after the second immunization. All data from the mPlex-Flu, HAI, and MN assays were adjusted for dose difference using linear mixed effects models, as previously described (27, 28).

### mPLEX-Flu Analysis

We estimated the concentrations of anti-HA IgG antibodies against a 45 HA antigen panel of influenza viruses using the mPLEX-Flu assay, as described previously(25, 34). All influenza HA sequence identifiers uesd are listed in the TABLE S1 and the HA genetic distance (phylogenic tree) is shown in FIG S1 A. The panel recombinant HA proteins were expressed by baculovirus system and purified Ni^+^ affinity column selection as previously described (34) and verified (FIG S1 B.

The calculation of individual IgG concentrations for each influenza strain anti-HA IgG was performed using standard curves generated from five-parameter logistic regression models (27, 28). All IgG concentration results from the mPlex-Flu assay was adjusted using linear mixed effects models accounting for the group, day, and group-day interactions for each H5 vaccine strain. Covariates adjusted in the linear mixed effects models included age at enrollment, gender, ethnicity (Caucasian vs. non-Caucasian), dose (two dose levels: 15 and 90 *µg*), and analytic batch (five batches) factors (27, 28).

### Antigenic cartography of H5 influenza viruses generated by mPlex-Flu assay data

In order to estimate the antigenic distance of HA antigens of H5 influenza virus strains, we adopted the 17 H5 HA genes that covered all 10 clades/subclades strains of H5 from Dr. Paul Zhou from Institute Pasteur of Shanghai, Chinese Academy of Sciences, Shanghai, China (1). The 17 individual antisera against each H5 influenza virus strain were generated with mouse DNA vaccination as previously described (1), and shown in FIG S2 A. Using the mPlex-Flu assay, we evaluated the 17 anti-sera against a panel of 36 HA antigens to create a multi-dimensional matrix, after normalizing the dilution factors and subtracting the background levels, using generalized linear models with identity link functions (FIG S2 B). Classical multidimensional scaling was used to project multi-dimensional distances into two-dimensional antigenic cartography plots plots(30, 25). The coordinates for two-dimension antigenic cartography were further used to calculate the Euclidean distance between H5 influenza viruses to obtain the antigenic distance matrix(FIG S3).

### Relative antigenic landscapes of antibody response

Based on the antigenic distances generated above, and using the three vaccine strains as reference: A/Hong Kong/156/97 vaccine (HK97, clade 0) A/Vietnam/1203/04 (Vie04, clade 1) A/Indonesia/5/05 (Ind05, clade 2) a vaccine-strain relative antigenic distance matrix was selected. Next, relative antigenic antibody landscape-like figures were created by using the relative antigenic distance as the X-axis and the Y-axis is IgG antibody response. Data points were linked by LOWESS fit spline curves (Prism 8 software). A set of antibody response landscape-like plots were generated for each vaccination strain.

### H5 head and stalk specific antibody response

We used the mPlex-Flu assay to simultaneously assess the antibodies to the head and stalk domains of HA. We coupled Luminex beads with the head region of HA, which are purified recommbinant proteins of HA1 domain of H5/Ind05 and H9/A/guinea fowl/Hong Kong/WF10/1999 (gfHK99, H9). To detect the group 1 stalk-reactive antibodies, we used the chimeric cH5/H1 (head/stalk) and cH9/H1 proteins. For group 2 stalk-reactive antibodies, we used the cH5/H3 and cH7/H4 proteins kindly provided by Dr. Florian Krammer(52, 32, 34, 35).

### Reanalyses of HAI and MN data

Primary HAI and MN data were generated previously during the vaccine trial as described (24). Serum antibody responses to the homologous A/Indonesia/05/2005 PR8-IBCDC-RG2 virus were measured at the Southern Research Institute (6). We reanalyzed these data using linear mixed effects models, with correlations from repeated measurements within the same subject considered. The same predictors and covariates were used in the linear mixed effects models for the HAI and MN data analysis as for the mPLEX-Flu data analysis (27).

### Availability of data and materials

All data generated in this study are included in this published article and in the Supplementary Material.

## SUPPLEMENTAL MATERIAL

### Supplementary Material Main Text

**Supplementary Table 1:** The mPlex-Flu assay panel of seasonal influenza viruses, H5 clades and subclades.

**Supplementary Figure 1:** HA protein characters of 35 influenza virus A strains in mPlex-Flu assay. A. The phylogenetic tree was generated using HA amino acid sequences of the 35 influenza A virus strains obtained from the phylogenic tree maker on the Influenza Research Database Website (https://www.fludb.org/brc/home.spg?decorator=influenza). B.SDS-PAGE gel image of purified HA proteins of H5 influenza viral strains. C. HPLC analysis results of four representative HA proteins flowing through the Biosep-SEC-s4000 columns with the Bio-rad protein standards.

**Supplementary Figure 2:** Antigenic cartography is generated with a mouse DNA vaccination model. A. Mouse DNA vaccination strategy. B. Heat map of the multiple dimensional antibody data generated by the mPlex-Flu assay. Each mouse polyclonal antiserum was induced by DNA vaccination with a DNA plasmid encoding HA proteins, and the antibody levels in the sera were estimated by mPlex-Flu assay. C. Antigenic cartography of 36 influenza A strains assessed by mPlex-Flu assay with the Multiple Dimensional Scaling (MDS) method.

**Supplementary Figure 3:** The heat-map matrix of the antigenic distance between the 21 H5 influenza virus strains. The three vaccination strains are highlighted with red arrows.

**Supplementary Figure 4:** The correlation between the HA antibody response and HA antigenic similarity of A/Hong Kong/156/97 (HK97) to 21 H5 influenza virus strains. A. The HA antibody response landscape-like plots of each group using the relative HA antigenic distance of A/Hong Kong/156/97 (HK97, clade 0) as the reference strains (see material and methods). X-axis is relative antigenic distance; Y-axis is IgG antibody response; the spots were linked by LOWESS fit spline curve (Prism 8 software). B. The correlation of the HA antibody response to the HA-antigenic distance. The Δ change of antibody concentration of pre- and post-vaccination versus the relative HA antigenic distance of Vie04. The R squared values were calculated with simple linear regression analysis (Prism 8 software).

**Supplementary Figure 5:** The correlation between the HA antibody response and HA antigenic similarity between A/Vietnam/1203/2004 (Vie04) and 21 H5 influenza virus strains. A. The HA antibody response landscape-like plots of each group using the relative HA antigenic distance of A/Vietnam/1203/2004 (Vie04, clade 1) as the reference strains (see material and methods). X-axis is relative antigenic distance; Y-axis is IgG antibody response; the spots were linked by LOWESS fit spline curve (Prism 8 software). B. The correlation of the HA antibody response to the HA-antigenic distance. The Δ change of antibody concentration of pre- and post-vaccination versus the relative HA antigenic distance of Vie04. The R squared values were calculated with simple linear regression analysis (Prism 8 software).

**Supplementary Figure 6:** The correlation between the HA antibody response and HA antigenic similarity of A/Indonesia/5/05 (Ind05) to 21 H5 influenza virus strains.

A. The HA antibody response landscape-like plots of each group using the relative HA antigenic distance of A/Indonesia/5/05 (Ind05, clade 1) as the reference strains (see material and methods). X-axis is relative antigenic distance; Y-axis is IgG antibody response; the spots were linked by LOWESS fit spline curve (Prism 8 software). B. The correlation of the HA antibody response to the HA-antigenic distance. The Δ change of antibody concentration of pre- and post-vaccination verse the relative HA antigenic distance of Vie04. The R squared values were calculated with simple linear regression analysis (Prism 8 software).

**Supplementary Figure 7:** The IgG concentration of group 1 and 2 influenza virus strains was estimated by mPlex-Flu assay in the DMID 08-0059 study. The mPlex-Flu assay estimated the mean and standard deviation of IgG concentration for each group. Then the antibody concentrations were adjusted within the linear mixed-effects models, which included the following: age at enrollment, gender, ethnicity (Caucasian vs. non-Caucasian), dose (two dose levels: 15 and 90 *µg*), and batch (five batches). A. The mPlex-Flu assay estimated the antibody concentrations of group 1 influenza virus strains (including five human H1, one of each H2, H6, and H9). B. The antibody concentrations to group 2 influenza A virus strains (including four H3, and two H7 strains) were estimated by the mPlex-Flu assay.

**Supplementary Figure 8:** Prior vaccination with a monovalent influenza vaccine (MIV) increased the serum titers of hemagglutination-inhibition (HAI) and microneutralization (MN) antibody responses against three antigenically drifted virus vaccine strains, including new vaccine strain A/Indonesia/05/2005 (Ind05; clade 2), previous MIV strains A/Vietnam/1203/2004 (Vie04; clade 1), A/Hong Kong/156/1997 (HK97; clade 0). Naive subjects (Unprimed) received the MIV Ind05 strain and were subsequently boosted at day 28 with the same strain. A previous primed group, received the MIV Vie04 5 years prior, (Primed) then received a single dose of Ind05. The previous double primed MIV Vie04 and HK97 (Multiple). The mean and standard deviation of IgG concentration for each group were estimated by linear mixed effects models with group, day, and group-day interaction used to fit the data for each H5 vaccine strain. Covariates adjusted in the linear mixed effects models included the following: age at enrollment, gender, ethnicity (Caucasian vs. non-Caucasian), dose (two dose levels: 15 and 90 *µg*), and batch (five batches). * P<0.05, **P<0.01, ***P<0.001 Linear contrasts within the linear mixed effects models framework were used to do the statistical testing.

## ACKNOWLEDGMENTS

We would like to thank Dr. Paul Zhou from Institute Pasteur of Shanghai, Chinese Academy of Sciences, Shanghai, China for providing the H5 HA constructs used to generate the mouse anti-sera for antigenic cartography, and Dr. Florian Kramer, Ichan School of Medicine at Mount Sinai, New York, United States for several influenza single strain and chimeric HA constructs.

This work was supported by the National Institutes of Health Institute of Allergy, Immunology and Infectious Diseases grants R21 AI138500 (MZ, JW, AW, SP), R01 AI129518 (MZ, SPH, JW, AW, SP) and the University of Rochester Clinical and Translational Science Award UL1 TR002001 from the National Center for Advancing Translational Sciences of the National Institutes of Health (JW, DL, MZ). The content is solely the responsibility of the authors and does not necessarily represent the official views of the National Institutes of Health.

J.W. and M.Z. conceived of the project, designed and oversaw the experiments the experiments and analysis, and wrote the paper. J.T. provided the data and samples from the prior DMID 08-0059 study, and contributed to the study design. D.L. performed the statistical analyses and modeling. J.W. S.P. and A.W performed the experiments. S.P.H. contributed to the experimental design and wrote the paper. M.S. contributed substantially to the design and analysis of the imprinting experiments. All authors read and approved the manuscript.

We declare no competing interests.

## REFERENCES

1. Zhou F, Wang G, Buchy P, Cai Z, Chen H, Chen Z, Cheng G, Wan XF, Deubel V, Zhou P. 2012. A triclade DNA vaccine designed on the basis of a comprehensive serologic study elicits neutralizing antibody responses against all clades and subclades of highly pathogenic avian influenza H5N1 viruses. J Virol 86 (12):6970–8. doi:10.1128/JVI.06930-11.

2. Gostic KM, Ambrose M, Worobey M, Lloyd-Smith JO. 2016. Potent protection against H5N1 and H7N9 influenza via childhood hemagglutinin imprinting. Sci 354 (6313):722–726. doi:10.1126/science.aag1322.

3. Monto AS. 2005. The threat of an avian influenza pandemic. N Engl J Med 352 (4):323–5. doi:10.1056/NEJMp048343.

4. for Disease Control PC. December 12,2018 2018. CDC, (ed), Highly Pathogenic Asian Avian Influenza A(H5N1) Virus. Center for Disease Control https://www.cdc.gov/flu/avianflu/h5n1-virus.htm.

5. Organization WH. 2015. WHO, (ed), Influenza at the human-animal interface (Summary and assessment as of 6 January 2015). WHO https://www.who.int/influenza/human_animal_interface/Influenza_Summary_IRA_6

6. Treanor JJ, Wilkinson BE, Masseoud F, Hu-Primmer J, Battaglia R, O’Brien D, Wolff M, Rabinovich G, Blackwelder W, Katz JM. 2001. Safety and immunogenicity of a recombinant hemagglutinin vaccine for H5 influenza in humans. Vaccine 19 (13-14):1732–7. https://www.ncbi.nlm.nih.gov/pubmed/11166898.

7. Patel SM, Atmar RL, El Sahly HM, Guo K, Hill H, Keitel WA. 2012. Direct comparison of an inactivated subvirion influenza A virus sub-type H5N1 vaccine administered by the intradermal and intramuscular routes. J Infect Dis 206 (7):1069–77. doi:10.1093/infdis/jis402.

8. Subbarao K. 2018. Avian influenza H7N9 viruses: a rare second warning. Cell Res 28 (1):1–2. doi:10.1038/cr.2017.154.

9. Couch RB, Patel SM, Wade-Bowers CL, Nino D. 2012. A randomized clinical trial of an inactivated avian influenza A (H7N7) vaccine. PLoS One 7 (12):e49704. doi:10.1371/journal.pone.0049704.

10. Couch RB, Decker WK, Utama B, Atmar RL, Nino D, Feng JQ, Halpert MM, Air GM. 2012. Evaluations for in vitro correlates of immunogenicity of inactivated influenza a H5, H7 and H9 vaccines in humans. PLoS One 7 (12):e50830. doi:10.1371/journal.pone.0050830.

11. Goji NA, Nolan C, Hill H, Wolff M, Noah DL, Williams TB, Rowe T, Treanor JJ. 2008. Immune responses of healthy subjects to a single dose of intramuscular inactivated influenza A/Vietnam/1203/2004 (H5N1) vaccine after priming with an antigenic variant. J Infect Dis 198 (5):635–41. doi:10.1086/590916.

12. Baer J, Santiago F, Yang H, Wu H, Holden-Wiltse J, Treanor J, Topham DJ. 2010. B cell responses to H5 influenza HA in human subjects vaccinated with a drifted variant. Vaccine 28 (4):907–15. doi:10.1016/j.vaccine.2009.11.002.

13. Krammer F, Srivastava K, Alshammary H, Amoako AA, Awawda MH, Beach KF, Bermudez-Gonzalez MC, Bielak DA, Carreno JM, Chernet RL, Eaker LQ, Ferreri ED, Floda DL, Gleason CR, Hamburger JZ, Jiang K, Kleiner G, Jurczyszak D, Matthews JC, Mendez WA, Nabeel I, Mulder LCF, Raskin A J, Russo KT, Salimbangon AT, Saksena M, Shin AS, Singh G, Sominsky LA, Stadlbauer D, Wajnberg A, Simon V. 2021. Antibody Responses in Seropositive Persons after a Single Dose of SARS-CoV-2 mRNA Vaccine. N Engl J Med doi:10.1056/NEJMc2101667.

14. Nachbagauer R, Palese P. 2018. Development of next generation hemagglutinin-based broadly protective influenza virus vaccines. Curr Opin Immunol 53:51–57. doi:10.1016/j.coi.2018.04.001.

15. Francis J T. 1960. On the doctrine of original antigenic sin. Proc Am Philos Soc 1960 104:572–578.

16. Francis J T, Davenport FM, Hennessy AV. 1953. A serological recapitulation of human infection with different strains of influenza virus. Trans Assoc Am Physicians 66:231–9. https://www.ncbi.nlm.nih.gov/pubmed/13136267.

17. Cobey S, Hensley SE. 2017. Immune history and influenza virus susceptibility. Curr Opin Virol 22:105–111. doi:10.1016/j.coviro.2016.12.004.

18. Zhang A, Stacey HD, Mullarkey CE, Miller MS. 2019. Original Antigenic Sin: How First Exposure Shapes Lifelong Anti-Influenza Virus Immune Responses. J Immunol 202 (2):335–340. doi:10.4049/jim-munol.1801149.

19. Monto AS, Malosh RE, Petrie JG, Martin ET. 2017. The Doctrine of Original Antigenic Sin: Separating Good From Evil. J Infect Dis 215 (12):1782–1788. doi:10.1093/infdis/jix173.

20. Nachbagauer R, Choi A, Hirsh A, Margine I, Iida S, Barrera A, Ferres M, Albrecht RA, Garcia-Sastre A, Bouvier NM, Ito K, Medina RA, Palese P, Krammer F. 2017. Defining the antibody cross-reactome directed against the influenza virus surface glycoproteins. Nat Immunol 18 (4):464–473. doi:10.1038/ni.3684.

21. Krammer F, Palese P. 2013. Influenza virus hemagglutinin stalk-based antibodies and vaccines. Curr Opin Virol 3 (5):521–30. doi:10.1016/j.coviro.2013.07.007.

22. Who O, et al.. 2008. Toward a unified nomenclature system for highly pathogenic avian influenza virus (H5N1). Emerg infectious diseases 14 (7):e1.

23. Writing Committee of the Second World Health Organization Consultation on Clinical Aspects of Human Infection with Avian Influenza AV, Abdel-Ghafar AN, Chotpitayasunondh T, Gao Z, Hayden FG, Nguyen DH, de Jong MD, Naghdaliyev A, Peiris JS, Shindo N, Soeroso S, Uyeki TM. 2008. Update on avian influenza A (H5N1) virus infection in humans. N Engl J Med 358 (3):261–73. doi:10.1056/NEJMra0707279.

24. Nayak JL, Richards KA, Yang H, Treanor JJ, Sant A J. 2015. Effect of influenza A(H5N1) vaccine prepandemic priming on CD4+ T-cell re-sponses. J Infect Dis 211 (9):1408–17. doi:10.1093/infdis/jiu616.

25. Wang J, Hilchey SP, Hyrien O, Huertas N, Perry S, Ramanunninair M, Bucher D, Zand MS. 2015. Multi-Dimensional Measurement of Antibody-Mediated Heterosubtypic Immunity to Influenza. PLoS One 10 (6):e0129858. doi:10.1371/journal.pone.0129858.

26. Fonville JM, Wilks SH, James SL, Fox A, Ventresca M, Aban M, Xue L, Jones TC, L. NM, Pham QT, Tran ND, Wong Y, Mosterin A, Katzelnick LC, Labonte D, Le TT, van der Net G, Skepner E, Russell CA, Kaplan TD, Rimmelzwaan GF, Masurel N, de Jong JC, Palache A, Beyer WE, L. QM, Nguyen TH, Wertheim HF, Hurt AC, Osterhaus AD, Barr IG, Fouchier RA, Horby PW, Smith DJ. 2014. Antibody landscapes after influenza virus infection or vaccination. Sci 346 (6212):996–1000. doi:10.1126/science.1256427.

27. Li D, Wang J, Garigen J, Treanor JJ, Zand MS. 2019. Continuous Read-out versus Titer-Based Assays of Influenza Vaccine Trials: Sensitivity, Specificity, and False Discovery Rates. Comput Math Methods Med 2019:9287120. doi:10.1155/2019/9287120.

28. Li D, Wang J, Treanor JJ, Zand MS. 2019. Improved Specificity and False Discovery Rates for Multiplex Analysis of Changes in Strain-Specific Anti-Influenza IgG. Comput Math Methods Med 2019:3053869. doi:10.1155/2019/3053869.

29. Fonville JM, Wilks SH, James SL, Fox A, Ventresca M, Aban M, Xue L, Jones TC, L. NMH, Pham QT, Tran ND, Wong Y, Mosterin A, Katzelnick LC, Labonte D, Le TT, van der Net G, Skepner E, Russell CA, Kaplan TD, Rimmelzwaan GF, Masurel N, de Jong JC, Palache A, Beyer WEP, L. QM, Nguyen TH, Wertheim HFL, Hurt AC, Osterhaus A, Barr IG, Fouchier RAM, Horby PW, Smith DJ. 2014. Antibody landscapes after influenza virus infection or vaccination. Sci 346 (6212):996–1000. doi:10.1126/science.1256427.

30. Zand MS, Wang J, Hilchey S. 2015. Graphical representation of proximity measures for multidimensional data: Classical and metric multi-dimensional scaling. Math J 17. doi:10.3888/tmj.17-7.

31. Smith DJ, Lapedes AS, de Jong JC, Bestebroer TM, Rimmelzwaan GF, Osterhaus AD, Fouchier RA. 2004. Mapping the antigenic and genetic evolution of influenza virus. Sci 305 (5682):371–6. doi:10.1126/science.1097211.

32. Pica N, Hai R, Krammer F, Wang TT, Maamary J, Eggink D, Tan GS, Krause JC, Moran T, Stein CR, Banach D, Wrammert J, Belshe RB, Garcia-Sastre A, Palese P. 2012. Hemagglutinin stalk antibodies elicited by the 2009 pandemic influenza virus as a mechanism for the extinction of seasonal H1N1 viruses. Proc Natl Acad Sci U S A 109 (7):2573–8. doi:10.1073/pnas.1200039109.

33. Wu NC, Wilson IA. 2018. Structural insights into the design of novel anti-influenza therapies. Nat Struct Mol Biol 25 (2):115–121. doi:10.1038/s41594-018-0025-9.

34. Wang J, Hilchey SP, DeDiego M, Perry S, Hyrien O, Nogales A, Garigen J, Amanat F, Huertas N, Krammer F, Martinez-Sobrido L, Topham DJ, Treanor JJ, Sangster MY, Zand MS. 2018. Broad cross-reactive IgG responses elicited by adjuvanted vaccination with recombinant influenza hemagglutinin (rHA) in ferrets and mice. PLoS One 13 (4):e0193680. doi:10.1371/journal.pone.0193680.

35. Tesini BL, Kanagaiah P, Wang J, Hahn M, Halliley JL, Chaves FA, Nguyen PQT, Nogales A, DeDiego ML, Anderson CS, Ellebedy AH, Strohmeier S, Krammer F, Yang H, Bandyopadhyay S, Ahmed R, Treanor JJ, Martinez-Sobrido L, Golding H, Khurana S, Zand MS, Topham DJ, Sangster MY. 2019. Broad Hemagglutinin-Specific Memory B Cell Expansion by Seasonal Influenza Virus Infection Reflects Early-Life Imprinting and Adaptation to the Infecting Virus. J Virol 93 (8). doi:10.1128/JVI.00169-19.

36. Ellebedy AH, Krammer F, Li GM, Miller MS, Chiu C, Wrammert J, Chang CY, Davis CW, McCausland M, Elbein R, Edupuganti S, Spearman P, Andrews SF, Wilson PC, Garcia-Sastre A, Mulligan MJ, Mehta AK, Palese P, Ahmed R. 2014. Induction of broadly cross-reactive antibody responses to the influenza HA stem region following H5N1 vaccination in humans. Proc Natl Acad Sci U S A 111 (36):13133–8. doi:10.1073/pnas.1414070111.

37. Wang J, Wiltse A, Zand MS. 2019. A Complex Dance: Measuring the Multidimensional Worlds of Influenza Virus Evolution and Anti-Influenza Immune Responses. Pathog 8 (4). doi:10.3390/pathogens8040238.

38. Belshe RB, Frey SE, Graham I, Mulligan MJ, Edupuganti S, Jackson LA, Wald A, Poland G, Jacobson R, Keyserling HL, Spearman P, Hill H, Wolff M, National Institute of A, Infectious Diseases-Funded V, Treatment Evaluation U. 2011. Safety and immunogenicity of influenza A H5 subunit vaccines: effect of vaccine schedule and antigenic variant. J Infect Dis 203 (5):666–73. doi:10.1093/infdis/jiq093.

39. Ledgerwood JE, Zephir K, Hu Z, Wei CJ, Chang L, Enama ME, Hendel CS, Sitar S, Bailer RT, Koup RA, Mascola JR, Nabel GJ, Graham BS, Team Vrcs. 2013. Prime-boost interval matters: a randomized phase 1 study to identify the minimum interval necessary to observe the H5 DNA influenza vaccine priming effect. J Infect Dis 208 (3):418–22. doi:10.1093/infdis/jit180.

40. Andrews SF, Chambers MJ, Schramm CA, Plyler J, Raab JE, Kanekiyo M, Gillespie RA, Ransier A, Darko S, Hu J, Chen X, Yassine HM, Boyington JC, Crank MC, Chen GL, Coates E, Mascola JR, Douek DC, Graham BS, Ledgerwood JE, McDermott AB. 2019. Activation Dynamics and Immunoglobulin Evolution of Preexisting and Newly Generated Human Memory B cell Responses to Influenza Hemagglutinin. Immun 51 (2):398–410 e5. doi:10.1016/j.immuni.2019.06.024.

41. Halliley JL, Khurana S, Krammer F, Fitzgerald T, Coyle EM, Chung KY, Baker SF, Yang H, Martinez-Sobrido L, Treanor JJ, Subbarao K, Golding H, Topham DJ, Sangster MY. 2015. High-Affinity H7 Head and Stalk Domain-Specific Antibody Responses to an Inactivated Influenza H7N7 Vaccine After Priming With Live Attenuated Influenza Vaccine. J Infect Dis 212 (8):1270–8. doi:10.1093/infdis/jiv210.

42. Galli G, Medini D, Borgogni E, Zedda L, Bardelli M, Malzone C, Nuti S, Tavarini S, Sammicheli C, Hilbert AK, Brauer V, Banzhoff A, Rappuoli R, Del Giudice G, Castellino F. 2009. Adjuvanted H5N1 vaccine induces early CD4+ T cell response that predicts long-term persistence of protective antibody levels. Proc Natl Acad Sci U S A 106 (10):3877–82. doi:10.1073/pnas.0813390106.

43. Khurana S, Coyle EM, Dimitrova M, Castellino F, Nicholson K, Del Giudice G, Golding H. 2014. Heterologous prime-boost vaccination with MF59-adjuvanted H5 vaccines promotes antibody affinity maturation towards the hemagglutinin HA1 domain and broad H5N1 cross-clade neutralization. PLoS One 9 (4):e95496. doi:10.1371/journal.pone.0095496.

44. Levine MZ, Holiday C, Jefferson S, Gross FL, Liu F, Li S, Friel D, Boutet P, Innis BL, Mallett CP, Tumpey TM, Stevens J, Katz JM. 2019. Heterologous prime-boost with A(H5N1) pandemic influenza vaccines induces broader cross-clade antibody responses than homologous prime-boost. NPJ Vaccines 4:22. doi:10.1038/s41541-019-0114-8.

45. Van Hoeven N, Fox CB, Granger B, Evers T, Joshi SW, Nana GI, Evans SC, Lin S, Liang H, Liang L, Nakajima R, Felgner PL, Bowen RA, Marlenee N, Hartwig A, Baldwin SL, Coler RN, Tomai M, Elvecrog J, Reed SG, Carter D. 2017. A Formulated TLR7/8 Agonist is a Flexible, Highly Potent and Effective Adjuvant for Pandemic Influenza Vaccines. Sci Rep 7:46426. doi:10.1038/srep46426.

46. Ellebedy AH, Nachbagauer R, Jackson KJL, Dai YN, Han J, Alsoussi WB, Davis CW, Stadlbauer D, Rouphael N, Chromikova V, McCausland M, Chang CY, Cortese M, Bower M, Chennareddy C, Schmitz A J, Zarnitsyna VI, Lai L, Rajabhathor A, Kazemian C, Antia R, Mulligan MJ, Ward AB, Fremont DH, Boyd SD, Pulendran B, Krammer F, Ahmed R. 2020. Adjuvanted H5N1 influenza vaccine enhances both cross-reactive memory B cell and strain-specific naive B cell responses in humans. Proc Natl Acad Sci U S A 117 (30):17957–17964. doi:10.1073/pnas.1906613117.

47. Ekiert DC, Kashyap AK, Steel J, Rubrum A, Bhabha G, Khayat R, Lee JH, Dillon MA, O’Neil RE, Faynboym AM, Horowitz M, Horowitz L, Ward AB, Palese P, Webby R, Lerner RA, Bhatt RR, Wilson IA. 2012. Cross-neutralization of influenza A viruses mediated by a single antibody loop. Nat 489 (7417):526–32. doi:10.1038/nature11414.

48. Krause JC, Tsibane T, Tumpey TM, Huffman CJ, Basler CF, Crowe J J E. 2011. A broadly neutralizing human monoclonal antibody that recognizes a conserved, novel epitope on the globular head of the influenza H1N1 virus hemagglutinin. J Virol 85 (20):10905–8. doi:10.1128/JVI.00700-11.

49. Whittle JR, Zhang R, Khurana S, King LR, Manischewitz J, Golding H, Dormitzer PR, Haynes BF, Walter EB, Moody MA, Kepler TB, Liao HX, Harrison SC. 2011. Broadly neutralizing human antibody that recognizes the receptor-binding pocket of influenza virus hemagglutinin. Proc Natl Acad Sci U S A 108 (34):14216–21. doi:10.1073/pnas.1111497108.

50. Ohshima N, Iba Y, Kubota-Koketsu R, Asano Y, Okuno Y, Kurosawa Y. 2011. Naturally occurring antibodies in humans can neutralize a variety of influenza virus strains, including H3, H1, H2, and H5. J Virol 85 (21):11048–57. doi:10.1128/JVI.05397-11.

51. Schmidt AG, Therkelsen MD, Stewart S, Kepler TB, Liao HX, Moody MA, Haynes BF, Harrison SC. 2015. Viral receptor-binding site antibodies with diverse germline origins. Cell 161 (5):1026–1034. doi:10.1016/j.cell.2015.04.028.

52. Hai R, Krammer F, Tan GS, Pica N, Eggink D, Maamary J, Margine I, Albrecht RA, Palese P. 2012. Influenza viruses expressing chimeric hemagglutinins: globular head and stalk domains derived from different subtypes. J Virol 86 (10):5774–81. doi:10.1128/JVI.00137-12.

